# A novel mitochondrial enriched antioxidant protects neurons against acute oxidative stress

**DOI:** 10.1101/109439

**Authors:** Nicola J. Drummond, Nick O. Davies, Janet E. Lovett, Mark R. Miller, Graeme Cook, Thomas Becker, Catherina G. Becker, Donald B. McPhail, Tilo Kunath

## Abstract

Excessive reactive oxygen species (ROS) can damage proteins, lipids, and DNA, which result in cell damage and death. The outcomes can be acute, as seen in stroke, or more chronic as observed in age-related diseases such as Parkinson’s disease. Here we investigate the antioxidant ability of a novel synthetic flavonoid, Proxison (7-decyl-3-hydroxy-2-(3,4,5-trihydroxyphenyl)-4-chromenone), using a range of *in vitro* and *in vivo* approaches. We show that, while it has radical scavenging ability on par with other flavonoids in a cell-free system, Proxison is orders of magnitude more potent than natural flavonoids at protecting neural cells against oxidative stress and is capable of rescuing damaged cells. The unique combination of a lipophilic hydrocarbon tail with a modified polyphenolic head group promotes efficient cellular uptake and mitochondrial localisation of Proxison. Importantly, *in vivo* administration of Proxison demonstrated effective and well tolerated neuroprotection against oxidative stress in a zebrafish model of dopaminergic neuronal loss.

## Introduction

Oxidative stress has been implicated in a wide range of age-related conditions, including heart disease, cancer, diabetes and neurodegenerative diseases^1-4^. In healthy cells there is a balance between the production of reactive oxygen species (ROS) and their removal by antioxidants, with discrete generation of ROS playing an essential role regulating cell signaling and function^5,6^. Excessive production of ROS results in oxidative stress causing aberrant cell signaling, damage to cellular components, and subsequent disease. A number of ROS are involved in cellular oxidative stress, including superoxide radicals, hydrogen peroxide and hydroxyl radicals. The chemical reactivity of each of these and their site of production within the cell determines their ability to damage cellular components^7,8^. Mitochondria produce superoxide radicals and hydrogen peroxide as by-products of ATP production. The major site of superoxide production was determined to be the flavin mononucleotide of complex I in the mitochondrial electron transport chain, but the coenzyme Q pool and the mitochondrial electron transport chain complex III are also capable of ROS production^9-12^. Acute increased superoxide production at complex I occurs in stroke through a conserved mechanism^13,14^. This results in oxidative damage and neuronal cell death by mechanisms that include increased H_2_O_2_ release^15,16^.

Chronic exposure to oxidative stress has been implicated in numerous age-related degenerative conditions, including Parkinson’s disease where age is the most significant risk factor. Post-mortem analysis of Parkinson’s patients brains showed evidence of oxidative damage and mitochondrial complex I dysfunction in the substantia nigra^3,17-20^. During earlier stages of Parkinson’s disease, patients also show evidence of mitochondrial dysfunction and oxidative damage in their platelets and plasma^21,22^. However, it is difficult to distinguish when in the disease process this damage occurs and how influential it is in the disease progression. The use of novel potent antioxidants in combination with refined disease models will be key to define the role for oxidative stress in disease pathogenesis, and to pinpoint when and where this occurs. If it proves to be an early direct defect, it opens up an exciting new field of potential therapeutic targets for intervention of age-related disease.

Flavonoids are a large group of polyphenolic antioxidant compounds found in plants that can directly scavenge ROS. Their antioxidant activity is a result of the efficiency by which they can donate hydrogen atoms from their multiple hydroxyl groups to free radicals, a mechanism that is facilitated by the extended conjugation afforded through the π-electron system of the core flavonoid molecular scaffold^23,24^. The dietary flavonoids, quercetin and myricetin are amongst the most potent^23-25^, although they are poorly absorbed from the diet^26-28^. Furthermore, their physicochemical attributes mitigate against effective uptake and distribution in the cell. However, cell absorption characteristics and bioactivity can be improved significantly by chemical modification of the parent myricetin through attachment of lipophilic alkyl chains of different lengths and by removal of hydroxyl groups that do not contribute to the antioxidant potential. A rational drug design approach, based on structure-activity relationships of natural and modified flavonoid antioxidants led to Proxison (7-decyl-3-hydroxy-2-(3,4,5-trihydroxyphenyl)-4-chromenone) as one of the most promising of these synthetic compounds^29^. This compound comprises of a straight chain C_10_ hydrocarbon tail covalently linked to a flavonoid head group similar to myricetin (Figure 1a). In terms of prevention of lipid peroxidation, Proxison exhibited superior protection when compared to structurally similar compounds^29^. Recently, Proxison was shown to reduce proliferation of a breast cancer cell line at high concentrations, but its antioxidant functions at physiological concentrations have not been investigated^30^.

**Figure 1.**
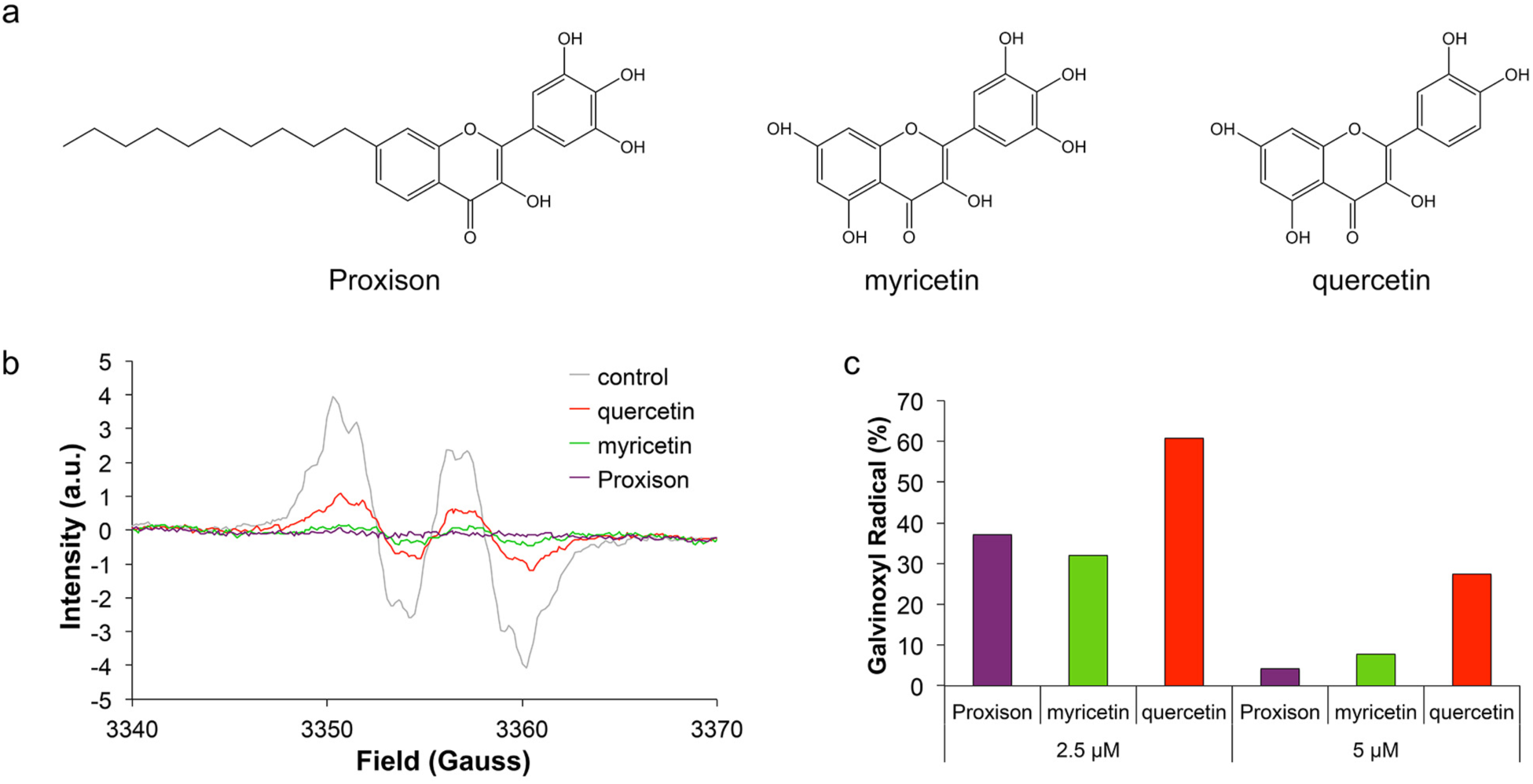
Chemical structures and radical scavenging ability of antioxidants. (a) The chemical structures of Proxison, myricetin, and quercetin. (b) Galvinoxyl EPR spectra in the absence or presence of 5 µM antioxidants. (c) The percentage of the galvinoxyl radical remaining after incubation with 2.5 µM or 5 µM of Proxison, myricetin and quercetin.

The aim of this study was to examine the mechanisms of antioxidant activity of Proxison using *in vitro* and *in vivo* model systems. This compound had potent radical scavenging capabilities in a cell-free system and was highly protective in a neuronal cell model of acute oxidative stress. *In vivo*, Proxison efficiently protected against 6-hydroxydopamine (6-OHDA) induced death of dopaminergic neurons in a zebrafish model. We demonstrate that Proxison is highly cell permeable and enriched in mitochondria, key characteristics for its protective antioxidant function in neurons. Therefore, Proxison represents a novel class of mitochondrial-targeted antioxidant, which is well tolerated and effective in promoting cell survival in conditions where oxidative stress is the major cause of cell death.

## Results

### Radical scavenging capabilities of flavonoid antioxidants

To directly compare the scavenging ability of Proxison to the structurally related natural flavonoids, myricetin and quercetin (Figure 1a), we used the stable free radical galvinoxyl in a cell-free assay. Galvinoxyl has one unpaired electron, which is extensively delocalised throughout the molecule and gives rise to a characteristic electron paramagnetic resonance (EPR) spectrum as a result of the interaction of the electron spin with the nuclear spins of protons in the molecule. All three flavonoids efficiently scavenged galvinoxyl radicals and mediated a concentration-dependent reduction in EPR signal intensity (Figure 1b,c). The scavenging ability of Proxison was equal to that of myricetin, and both were more than twice as potent as quercetin (Figure 1c).

### Protection against acute oxidative stress in cells

Next, we compared Proxison to myricetin, quercetin, and other antioxidants for their ability to protect against oxidative stress in a neural cell culture model, SH-SY5Y neuroblastoma cells. These cells were treated with the organic peroxide *tert*-butyl hydroperoxide (*t*BHP) to generate peroxyl radicals and cause acute oxidative stress^31^. Cells exhibited a significant reduction in cell metabolic activity (MTS tetrazolium reduction assay) after 5 h treatment with 200-800 µM *t*BHP (Supplementary Figure 1). 400 µM *t*BHP was found to be optimal as it elicited a robust and reproducible stressor effect in this cell-based assay.

Proxison used at 5 and 10 µM significantly prevented the reduction in cell metabolic activity caused by 400 µM *t*BHP treatment (Figure 2a). Quercetin also showed significant protection between 25-100 µM, however, a 10-fold higher concentration was required to exert a level of protection similar to Proxison. In contrast, myricetin showed no appreciable protection against *t*BHP even at 100 µM. Other antioxidants, including vitamin E, Idebenone (a synthetic derivative of coenzyme Q10) and coenzyme Q10 showed no protection in this assay. Furthermore, morphological signs consistent with lipid peroxidation, including membrane blebbing, which were observed by 5 h of *t*BHP treatment, were rescued by Proxison and high-dose quercetin, but not myricetin (Figure 2b).

**Figure 2.**
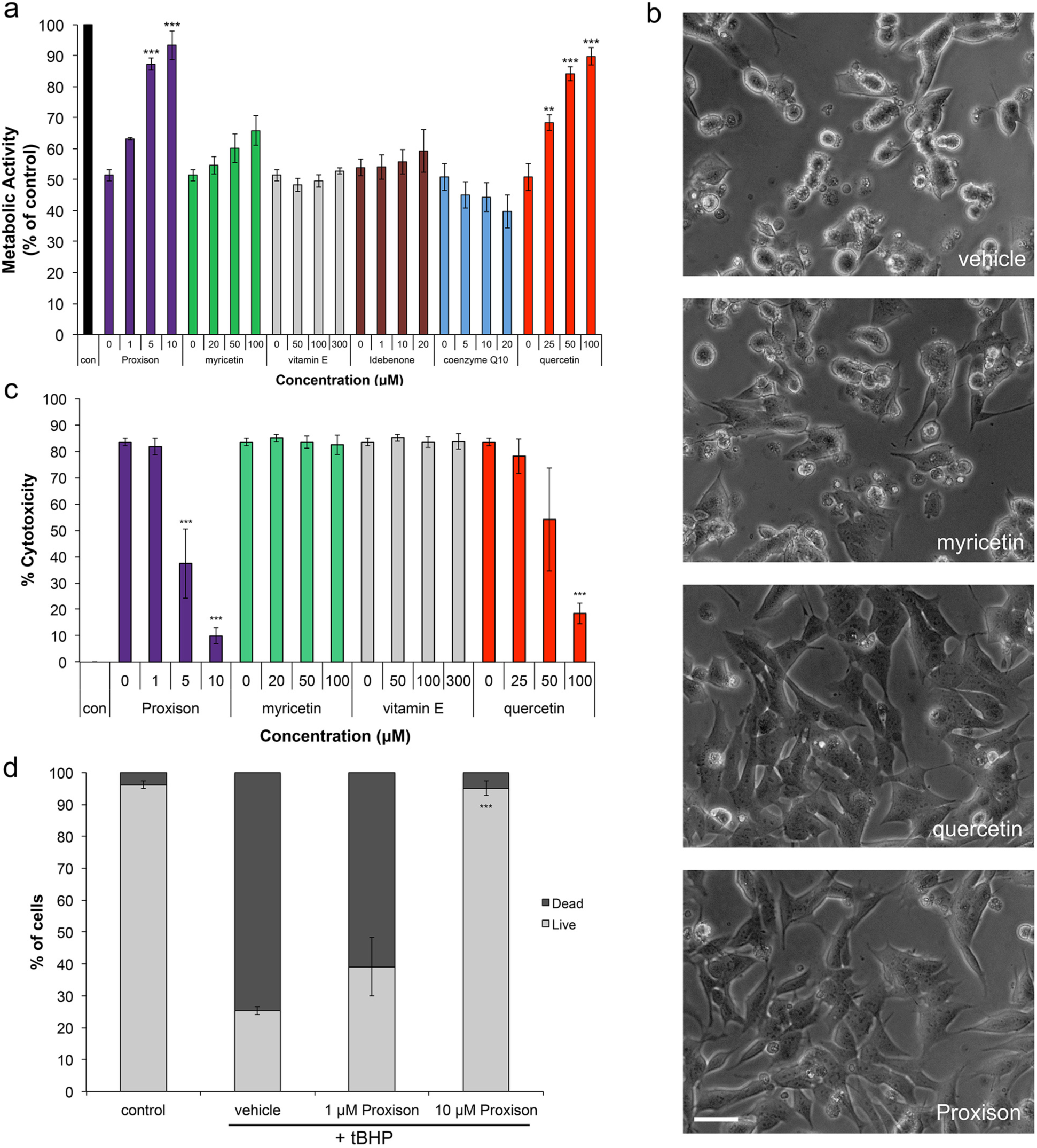
Antioxidant protection against acute oxidative stress. (a) Metabolic activity of SHSY5Y cells treated with 400 µM *t*BHP for 4 hours in the absence or presence of differing concentrations of 6 antioxidants (n=3). (b) Phase-contrast microscopy of *t*BHP-treated SH-SH5Y cells treated with DMSO (vehicle), myricetin, quercetin, or Proxison. Scale bar, 75 µm. (c) *t*BHP-induced cytotoxicity was determined at 16 hours with the LDH assay in the absence or presence of 4 antioxidants (n=3). (d) The percentage of dead cells was assessed at 16 hours by PI FACS analysis for Proxison-treated cells (n=2). ANOVA with a Tukey post-hoc test was performed to assess the significant difference between antioxidant-treated and vehicle-treated cells in (a), (c), and (d); ** p<0.01, *** p<0.001.

Oxidative stress can also result in cytotoxicity, so we next assessed cell death by the lactate dehydrogenase (LDH) assay. *t*BHP treatment of SH-SY5Y cells for 16 h caused a significant deterioration of cell membrane integrity resulting in leakage of cytoplasmic LDH into the media. Proxison (5-10 µM) and a high concentration of quercetin (100 µM) significantly prevented the cytotoxic effects of *t*BHP, while myricetin and vitamin E offered no protection (Figure 2c). Similarly, we further confirmed the cytoprotective capabilities of Proxison with a propidium iodide (PI) exclusion assay (Figure 2d). Less than 30% of *t*BHP-treated cells were viable after 16 hours as determined by this assay. However, pre-incubation with Proxison at 10 µM, but not 1 µM, protected against cell death to levels that were not significantly different from non-treated control cells (Figure 2d).

### Flavonoid antioxidant cellular uptake and intracellular localisation

Given that the antioxidants were similarly potent in cell-free assays, the differential protective properties of Proxison over myricetin and quercetin in cells could be due to differences in cellular uptake or subcellular localisation. To monitor intracellular uptake and localisation of the different compounds we took advantage of the endogenous fluorescence characteristics of these molecules^32,33^. Green fluorescence of Proxison, myricetin and quercetin was confirmed using a plate reader with excitation 485 nm and emission 520 nm (Supplementary Figure 2).

Cells were incubated with Proxison (10 µM), quercetin (100 µM), or myricetin (100 µM) for 2 h and washed out before processing for confocal imaging. Whereas quercetin accumulated in nuclei, Proxison strikingly localised to filamentous perinuclear structures exclusively in the cytoplasm (Figure 3a). The subcellular localisation of Proxison was confirmed to be enriched in mitochondria by co-localization with Mitotracker Deep Red (Figure 3b). Despite an emission profile similar to quercetin (Supplementary Figure 2), myricetin was not taken up into cells, as the fluorescence observed in treated cells was similar to the background in control cells (Figure 3a). Cellular uptake was further investigated by FACS analysis showing significant intracellular levels of Proxison and quercetin, whereas myricetin was undetectable (Figure 3c,d). This suggests the lack of protective activity of myricetin compared to Proxison and quercetin reflects its inability to enter the cell. Hence, Proxison was efficiently taken up by cells and accumulated in mitochondria.

**Figure 3.**
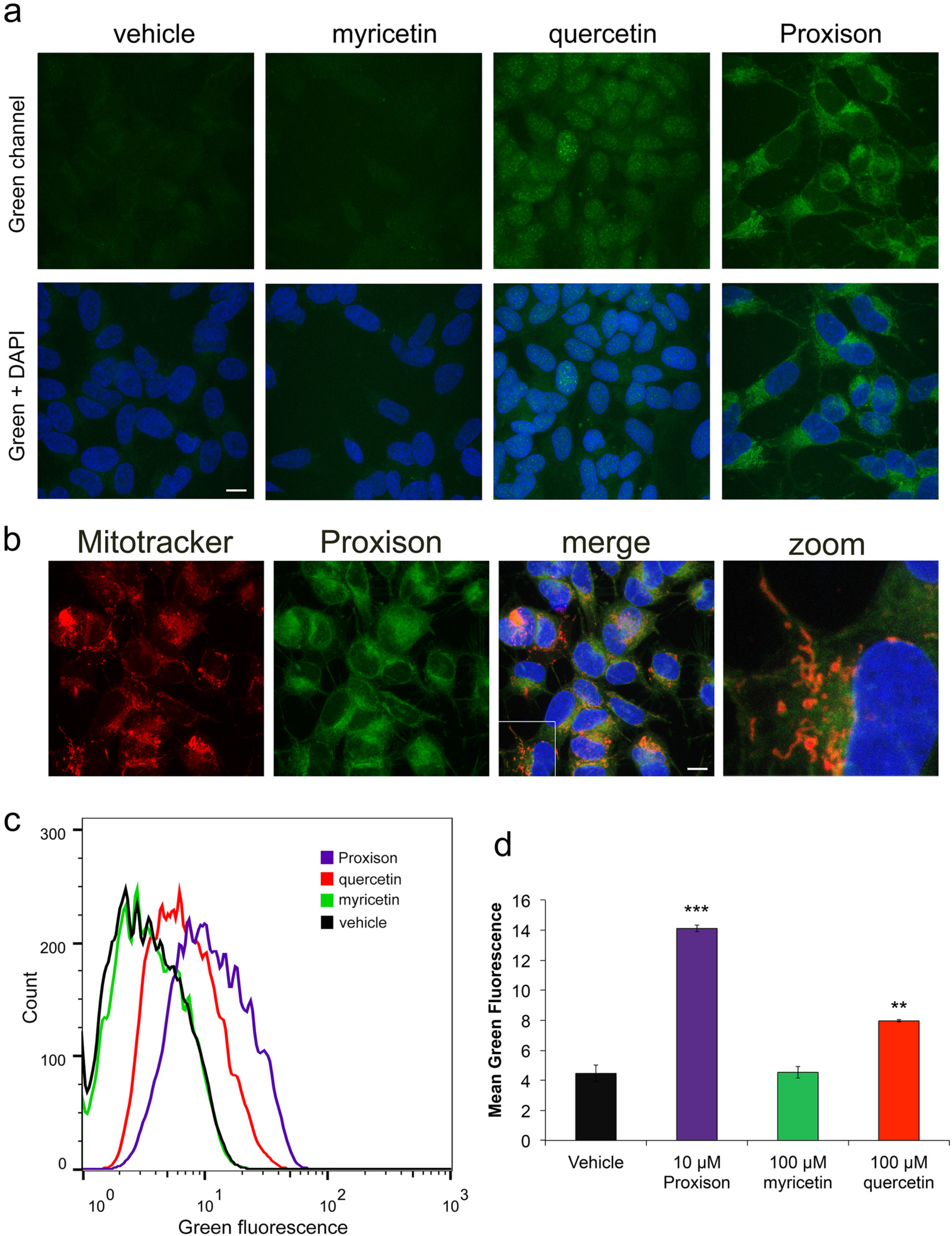
Cellular uptake and subcellular localisation of antioxidants. (a) Cellular uptake and localisation was observed in SH-SY5Y cells. Cells were incubated with Proxison, quercetin, myricetin or DMSO (Vehicle) before fixing, counterstaining with DAPI (blue), and confocal imaging, scale bar: 11 µm. (b) Mitochondrial localisation of Proxison was assessed by co-incubation with Mitotracker Deep Red prior to fixation and confocal imaging, scale bar: 11 µm. (c) Representative FACS plots of SH-SY5Y cells incubated with Proxison, quercetin, myricetin or DMSO (Vehicle) for 2 hours. (d) Mean green fluorescence by FACS of live cells incubated with each antioxidant (n=2). ANOVA with a Tukey post-hoc test was performed; ** p<0.01, *** p<0.001.

### Proxison prevents intracellular radical production and downstream effects of oxidative stress

To assess intracellular scavenging potency of Proxison, the ROS biosensor CellROX® Deep Red was used^34^. Using far-red fluorescence intensity as a read out of radical production, *t*BHP was found to significantly increase intracellular ROS in SH-SY5Y cells, and Proxison was capable of preventing this ROS production (Figure 4a). Quantification of far-red signals confirmed Proxison effectively blocked the accumulation of *t*BHP-generated radicals to the level of untreated cells (Figure 4b).

**Figure 4.**
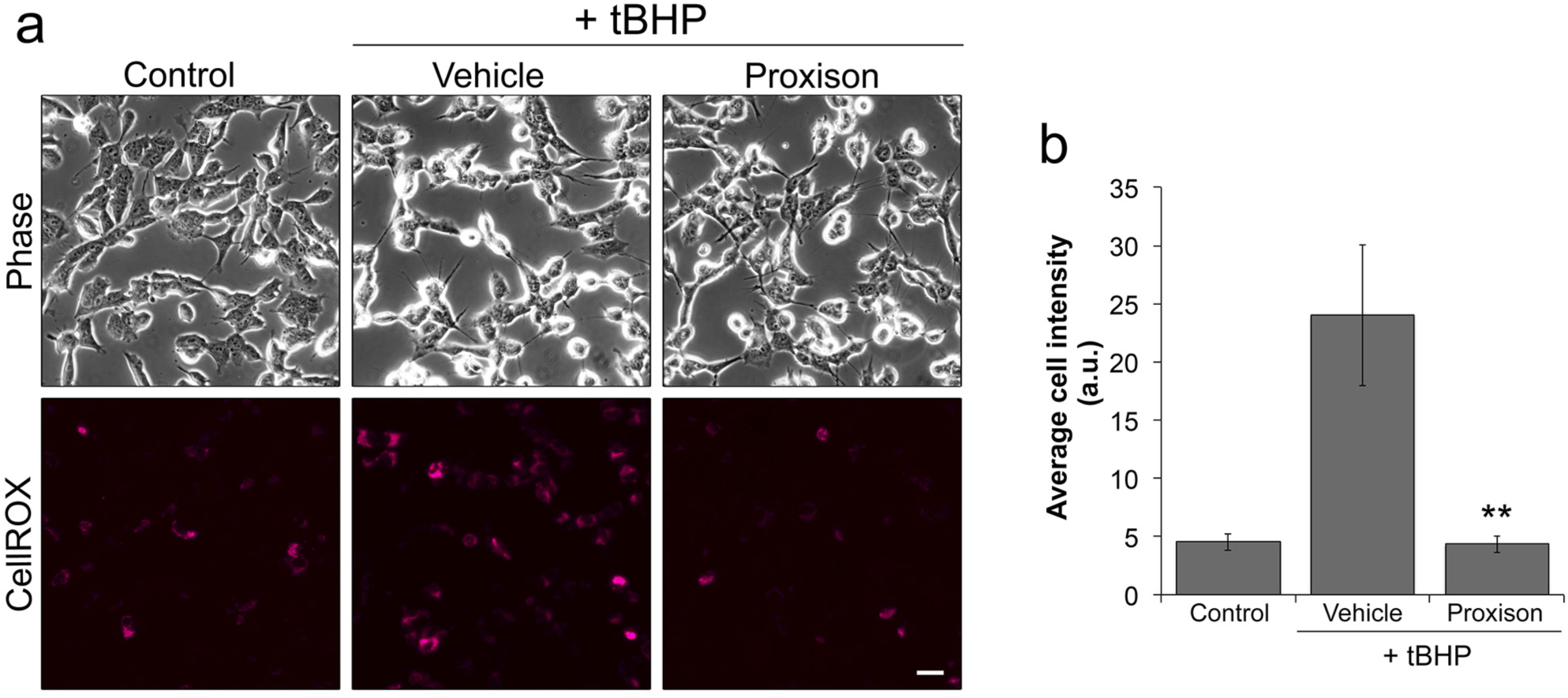
Intracellular scavenging of *t*BHP-induced radicals by antioxidants. (a) Intracellular ROS production in SH-SY5Y cells treated with *t*BHP in the absence or presence of Proxison was observed with CellROX® Deep Red. (b) The average intensity of CellROX® Deep Red (far red) fluorescence of each cell as quantified with Volocity. Bars represent the mean from four independent experiments with duplicate cover slips. Error bars represent the SEM. ANOVA with the post-hoc Tukey test was performed to assess significant difference to vehicle-treated cells; ** p<0.01.

A major mechanism in the cellular defense against oxidative stress involves activation of the Nrf2-Keap1 antioxidant response signaling pathway^35^. This requires nuclear localization of a key transcription factor NRF2 to activate a number of antioxidant related genes^35-37^. To determine whether Proxison is capable of preventing downstream consequences of *t*BHP induced oxidative stress we analysed subcellular localisation and action of NRF2. Nuclear NRF2 increased upon *t*BHP treatment, an effect that was significantly prevented in the presence of Proxison (Figure 5a,b). To confirm NRF2 was active in the nucleus, we quantified expression of two NRF2-regulated antioxidant response element (ARE) genes, *NQO1* and *HMOX1*. The increased transcription of both ARE genes caused by *t*BHP treatment was significantly attenuated by Proxison (Figure 5d). The transcript levels of *NRF2* itself were unaffected agreeing with the known post-translational mechanism of action for this transcription factor^37^.

**Figure 5.**
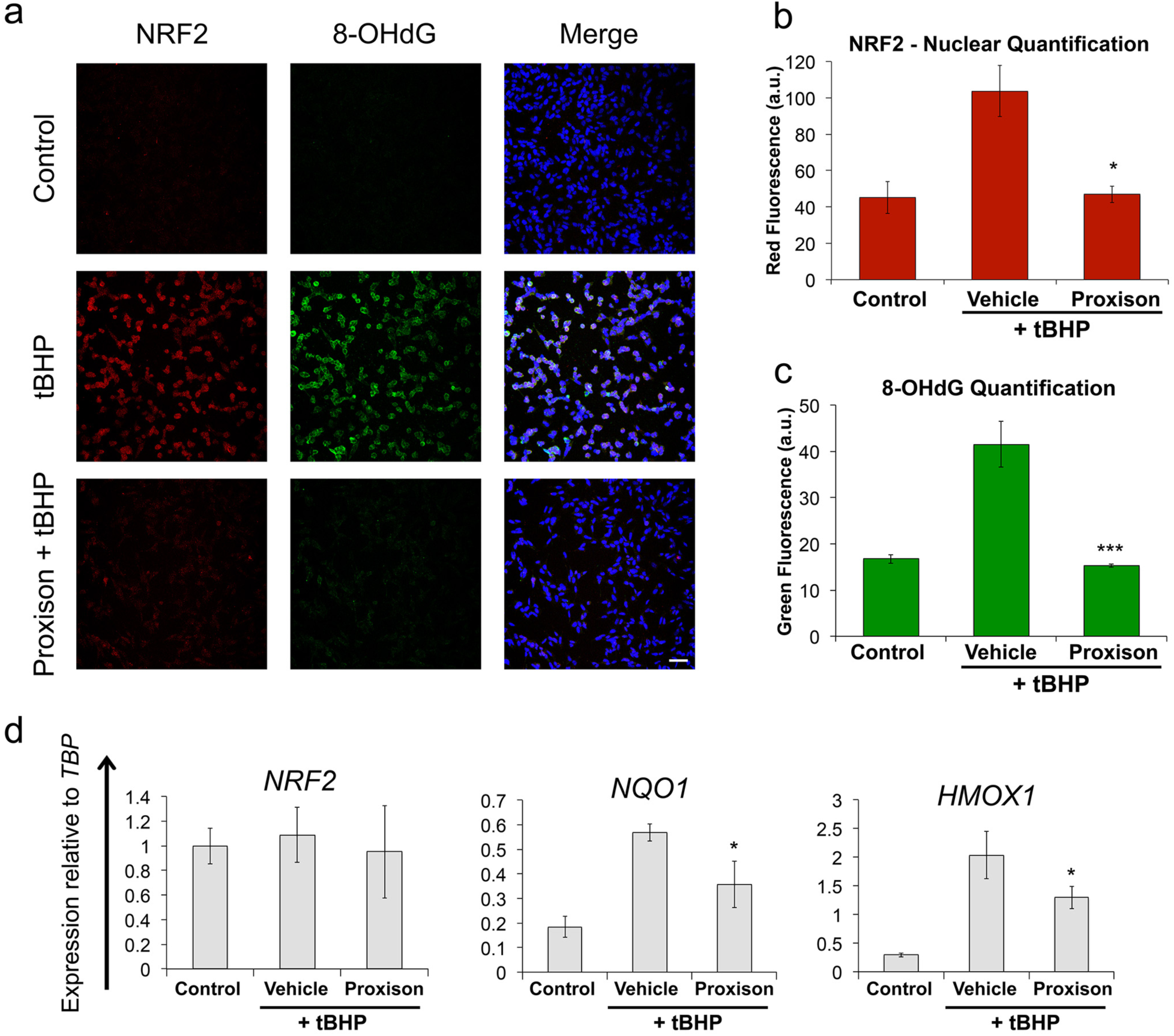
Proxison reduces *t*BHP-induced NRF2 oxidative stress response and prevents 8-OHdG DNA oxidative damage. (a) *t*BHP-treated SH-SY5Y cells in the absence or presence of Proxison were fixed and immunostained for NRF2 (red), 8-OHdG (green), and counterstained for the nuclear marker DAPI (blue). (b,c) The intensity of NRF2 red fluorescence (b) and 8-OHdG green fluorescence (c) in the nuclei as quantified using ImageJ. Bars in (b) and (c) represent the mean of three coverslips from independent experiments (n=3) with 2-4 images per coverslip, error bars represent the SEM. (d) RNA was extracted from similarly treated cells to measure gene expression of *NRF2*, *NQO1* and *HMOX1* relative to TATA-binding protein (*TBP*). Bars represent the mean of duplicate or triplicate wells (n=2-3). Error bars represent the standard deviation. ANOVA with the Tukey post-hoc test was performed in (b), (c) and (d) to assess significant difference to vehicle-treated cells; * p<0.05, *** p<0.001.

To determine whether Proxison protects against ROS-induced damage of DNA, we looked for the presence of 8-hydroxy-2'-deoxyguanosine (8-OHdG) in the nucleus. 8-OHdG is a major product of nucleic acid oxidation and its concentrations within a cell act as a read-out of oxidative stress induced damage^38^. Proxison was able to completely inhibit the accumulation of nuclear 8-OHdG caused by *t*BHP treatment (Figure 5a, c).

### Proxison protects dopaminergic neurons in a zebrafish model

To determine the ability of Proxison to protect neurons in an intact animal and to detect potential toxic side-effects, we assessed its ability to protect dopaminergic neurons in the brain of zebrafish embryos. The neurotoxic compound 6-hydroxydopamine (6-OHDA) was used to selectively induce death of dopaminergic and noradrenergic neurons in this model, as measured by counts of tyrosine hydroxylase (TH) positive neurons^39,40^. At 24 hours post fertilisation (hpf), zebrafish embryos were treated with 6-OHDA (250 µM for 24 h) in the absence or presence of the antioxidants Proxison or quercetin. At 48 hpf embryos were fixed and immunostained for TH prior to collection of z-stack images of each embryo head by confocal microscopy (Figure 6a), and the number of TH positive cells were quantified (Figure 6b). Embryos treated with 6-OHDA consistently had 30% less TH positive cells than control embryos, while co-incubation with Proxison or quercetin showed significant protection against 6-OHDA-induced neuronal cell loss (Figure 6a,b). None of the treatments influenced survival or normal development of the embryos, showing that Proxison was not toxic and was well tolerated at the concentrations used that elicited neuroprotective effects *in vivo*.

**Figure 6.**
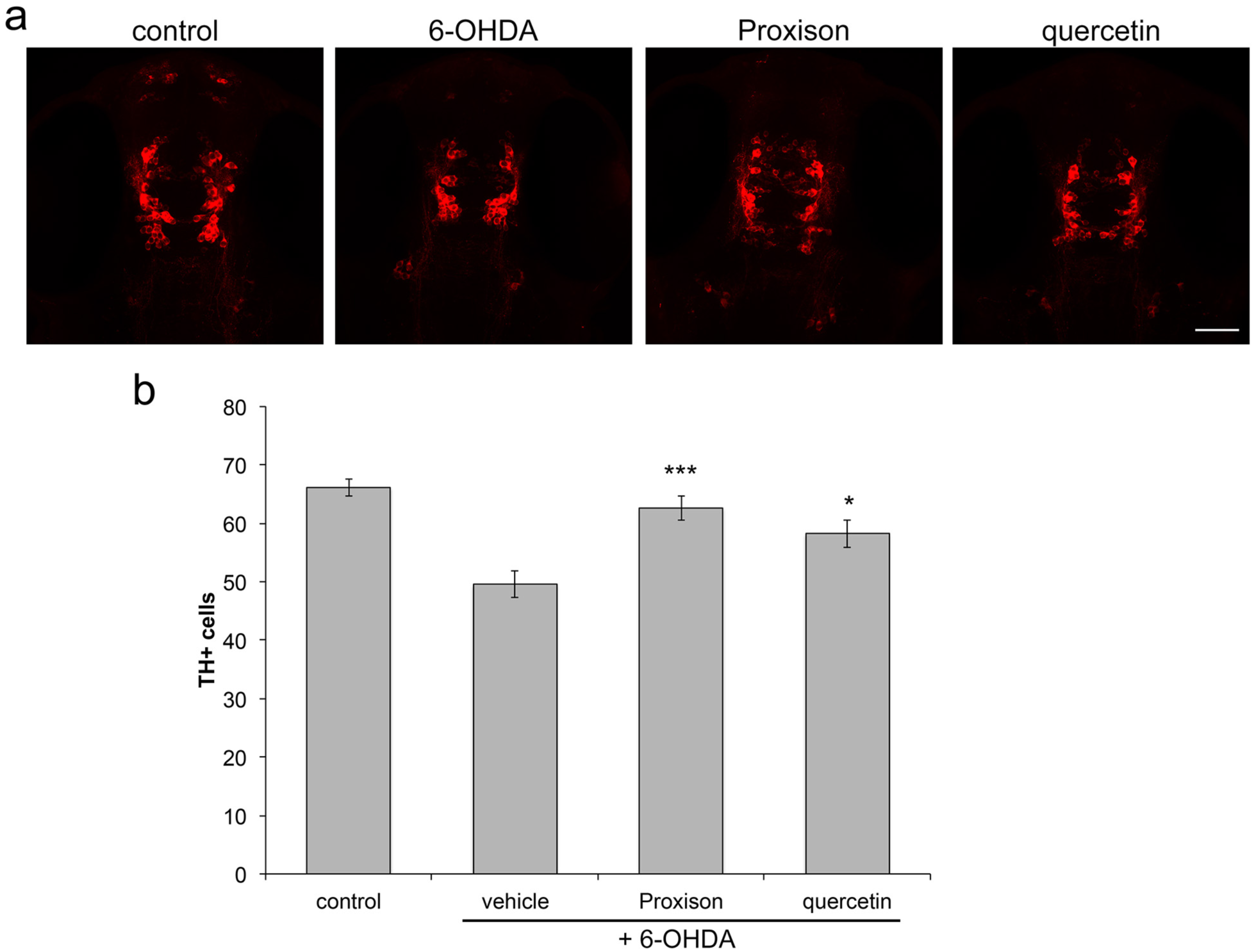
Antioxidant protection against 6-OHDA-induced dopaminergic cell death in zebrafish embryos. (a) Zebrafish embryos at 24 hours post-fertilisation (hpf) were treated with 6-OHDA for 24 hours in the presence or absence of the antioxidants Proxison or quercetin (10 µM). Embryos were then fixed, immunostained for TH (red) and confocal images were taken of the embryo head viewed from the top, scale bar: 50 µm. (b) The number of TH-positive cells in each embryo was counted for each treatment group (n=24-30). Bars represent the mean number of TH-positive cells and error bars show the SEM. ANOVA with a Tukey post-hoc test was performed to assess the significant difference to the 6-OHDA treated embryos; * p<0.05, *** p<0.001.

## Discussion

Here we characterise the antioxidant profile of a novel synthetic flavonoid, Proxison. Although the intrinsic radical scavenging activity of Proxison is similar to its natural counterparts, it possesses a superior ability to protect against acute oxidative stress in cellular contexts. The lipophilic tail of Proxison enhances its cellular uptake and may improve its stability^29^. The subcellular localization of Proxison, enriching in the mitochondria, is also likely to be an important factor in its efficacy. Although myricetin was as potent as Proxison in a cell-free antioxidant assay, it was poorly taken up by cells, possibly due to its instability at neutral pH or its low lipophilicity^41,42^. Quercetin is readily taken up by cells, but was a less potent antioxidant scavenger. It accumulates in the nucleus for, as yet, unknown reasons^32^. This localization could explain why a 10-fold higher concentration of quercetin is needed to provide a similar level of protection as Proxison against *t*BHP-induced toxicity. The superior potency of Proxison may also be due to the phenoxyl radical (the oxidized form of Proxison) being recycled by intracellular antioxidants, or other reductive pathways, back to Proxison, as occurs for vitamin E^43^. Flavonoid antioxidants are also capable of chelating transition metal ions such as iron or copper thereby limiting the breakdown of *t*BHP to its initiating free radical products or inhibiting lipid peroxidation through the Fenton reaction^24^. Proxison may therefore be able to exert a protective effect by a mechanism other than direct scavenging of free radicals by hydrogen atom electron transfer reactions.

We demonstrated that Proxison scavenges intracellular ROS and prevents downstream oxidative damage to both membranes and DNA resulting from *t*BHP treatment. Induction of the NRF2 antioxidant response pathway was significantly dampened by Proxison, but not eliminated. Therefore, Proxison can act in conjunction with other cellular antioxidant responses to acute oxidative stress to promote cell survival. Importantly, the studies in zebrafish embryos showed that Proxison is also capable of protecting dopaminergic neurons against acute oxidative stress in an intact animal.

There are currently two published classes of synthetic mitochondrial-targeted antioxidants, one is the Mito compounds and the other is a group of small peptides known as the SS peptides. The Mito compounds contain a triphenylphosphonium ion (TPP), which has a positive charge attracting it to the mitochondrial matrix. The TPP ion has been attached to a number of known antioxidants, including vitamin E and coenzyme Q10^44,45^. MitoQ compounds are taken up into cells and accumulate in mitochondria only when the membrane potential is intact, which may not always be the case in damaged cells^44-46^. MitoQ was tested in a phase II clinical trial for symptomatic Parkinson’s disease patients. However, there was no change in the rate of disease progression^47^. Currently MitoQ is due to undergo a phase II clinical trial for chronic kidney disease (ClinicalTrials.gov Identifier: NCT02364648). SS peptides are short positively charged peptides that are cell permeable and localise to mitochondria^48,49^. SS peptide localisation is due to their lipophilicity and positive charge and is partially reliant on an intact mitochondrial membrane potential^49,50^. The lead SS peptide, SS-31 (Bendavia; MTP-131), is protective in rodent models of amyotrophic lateral sclerosis and myocardial infarction^51,52^. Bendavia is currently in clinical trials for heart failure (ClinicalTrials.gov Identifier: NCT02388464) and genetically confirmed mitochondrial disease (ClinicalTrials.gov Identifier: NCT02805790). Proxison defines a new class of mitochondrial-targeted antioxidant that becomes enriched in this organelle as a likely consequence of the lipophilicity of a long hydrocarbon tail combined with the unique redox properties of the polyphenolic head group^53^.

This novel antioxidant can be applied to investigate mitochondrial oxidative stress in disease models, like α-synucleinopathies and other neurodegeneration models. This would provide insight into disease processes and aid in development of novel treatments. In addition, Proxison could have applications for regenerative medicine where oxidative stress has been implicated in poor cell survival of transplanted cells, with the advantage that the molecule can be pre-loaded into cells prior to transplantation. Proxison could also have applications for conditions, such as stroke or cardiac infarction, in which a temporary, but acute, exposure to oxidative stress is experienced, as well as diseases in which oxidative stress and mitochondrial dysfunction are core features.

## Methods

### Galvinoxyl Electron Paramagnetic Resonance Spectroscopy

Galvinoxyl (182 µM, Sigma) was mixed with 2.5 µM or 5 µM Proxison (7-decyl-3-hydroxy-2-(3,4,5-trihydroxyphenyl)-4-chromenone, Antoxis), myricetin (Sigma) or quercetin (Sigma) in DMSO. Five minutes after addition measurements were taken using the Magnettech Miniscope MS200 EPR machine. Settings: centre field= 3364.05 G; Sweep= 69.59 G; Sweep Time= 20 s; Smooth= 0 s; Steps= 4096; Number of passes= 1 pass; Modulation= 1.5 G; Power= 7dB; Gain= 1E2. The percentage radical scavenging ability was calculated from the EPR signal intensity with antioxidant relative to the signal intensity of galvinoxyl alone.

### Cell Culture

The SH-SY5Y cell line was a kind gift from Professor David Porteous (Centre for Genomic and Experimental Medicine, University of Edinburgh). They were cultured at 37°C in a humidified atmosphere of 5% CO_2_ in DMEM supplemented with 10% FBS, 2 mM glutamine and 1 mM sodium pyruvate (all from Life Technologies). When cells were 70-80% confluent they were passaged using trypsin (Life Technologies).

### MTS Assay

The MTS assay was performed using the Promega CellTiter 96® Aqueous MTS Reagent Powder (Promega) with phenazine methosulphate (PMS; Sigma) as the electron coupler. SH-SY5Y cells were seeded at 1.25×10^5^ cells/cm^2^ in 96-well plates and incubated for 24 hours when either *t*BHP (Sigma) was added or cells were pre-incubated with antioxidants Proxison, myricetin, quercetin, vitamin E (Sigma), Idebenone (Antoxis) or coenzyme Q10 (Sigma) for 30 minutes before *t*BHP (400 µM) addition. Five hours later phase-contrast images were collected with an Olympus IX51 inverted microscope and the MTS assay was performed, according to manufacturer’s instructions. Briefly, 100 µl of PMS (0.92 mg/ml) was added to 2 ml of MTS (2 mg/ml), and then 20 µl of the MTS/PMS mixture was added to each well of a 96-well plate containing 100 µl of conditioned media. The plate was incubated at 37°C/5% CO_2_ for 1 hour when the absorbance at 490 nm was measured using the FLUOstar OMEGA microplate reader (BMG LabTech). The background absorbance from the media incubated with MTS/PMS was subtracted from the well absorbance to give the sample absorbance. The percentage of the sample absorbance relative to control treated cells absorbance gave the % metabolic activity.

### LDH Assay

The LDH-Cytotoxicity Assay Kit II (Source Bioscience) was used to quantify cell death due to *t*BHP treatment. SH-SY5Y cells were pre-incubated with antioxidants for 30 minutes when 400 µM *t*BHP was added to the antioxidants for 16 hours. Thirty minutes before the end of *t*BHP treatment cell lysis buffer was added to the high control wells. At the end of *t*BHP treatment the 96-well plate was gently mixed to distribute the LDH in the media. The plate was then centrifuged at 600g for 10 minutes to precipitate cells and debris. 10 µl of media from each well was moved to a new 96-well plate and 100 µl of LDH reaction mix was added to each well. The plate was then incubated at room temperature for 30-60 minutes when absorbance was read using the FLUOstar OMEGA microplate reader. The 650 nm reference absorbance was subtracted from the 450 nm absorbance reading to give the well absorbance. The background absorbance was subtracted from the well absorbance to give the sample absorbance. The sample absorbance was subtracted from control-cultured cells (low control) and divided by the cells that had been lysed (high control) minus the control cells (low control) to give the % cytotoxicity.

### PI FACS

The propidium iodide (PI) assay was performed to assess Proxison protection against *t*BHP-induced toxicity. SH-SY5Y cells were pre-incubated with Proxison (1 or 10 µM) for 30 minutes when 400 µM *t*BHP was added for 16 hours. Then the media was collected and cells were lifted using trypsin, washed in PBS with 2% FBS and resuspended in 100 µl PBS with 2% FBS. This cell solution was filtered and incubated with PI (1 µg/ml) at room temperature for 10 minutes. Then 300 µl PBS with 2% FBS was added to dilute the cells. Cells were kept on ice until FACS analysis data was collected using the FACS Calibur and post-acquisition analysis was performed using FlowJo X. The unstained sample was used to create gates to determine PI positive cells and the percentage of PI positive and negative cells was calculated.

### Antioxidant Fluorescence

Proxison, myricetin and quercetin (1 mM) in DMSO were added to a 96-well plate in triplicate. The intrinsic fluorescence of each compound was measured using a FLUOstar OMEGA microplate reader (BMG LabTech) with excitation 485 nm and emission 520 nm. DMSO background fluorescence was subtracted.

### Antioxidant Subcellular Localisation

SH-SY5Y cells were seeded on coverslips and loaded with the following antioxidants for 2 hours; 10 µM Proxison, 100 µM quercetin, 100 µM myricetin or DMSO as a vehicle control. Then cells were fixed in 4% PFA (Sigma) and counterstained with 4',6-diamidino-2-phenylindole (DAPI; Life Technologies) for 10 minutes. Coverslips were washed in MilliQ water and mounted onto slides using Vectashield (Vector Laboratories, H1000). Images were taken using the 63x objective of the Leica TCS SPE confocal microscope. The 488 nm laser was used to excite the antioxidants and emission was measured between 496-595 nm. To observe Proxison subcellular localization, SHSY5Y cells on coverslips (as above) were loaded with 100 nM Mitotracker Deep Red (Life Technologies) for 20 minutes before the addition of 10 µM Proxison for 2 hours. The cells were then fixed in 4% PFA and processed as described above. The 635 nm laser was used to excite Mitotracker Deep Red.

### Intracellular Antioxidant Fluorescence (FACS)

SH-SY5Y cells were treated as above for antioxidant localisation but 2 hours after antioxidant addition cells were lifted using trypsin, washed in PBS with 2% FBS and resuspended in 100 µl PBS with 2% FBS. The cell solution was then filtered and cells were incubated with PI (1 µg/ml) on ice. 300 µl PBS with 2% FBS was added. Cells were kept on ice until fluorescence was analysed using the FACS Calibur. Post-acquisition analysis using FlowJo X was used to exclude PI positive cells and to graph the cells positive for green (FL-1) fluorescence. Mean green fluorescence was calculated with FlowJo X.

### CellROX® Deep Red Assay

SH-SY5Y cells were seeded at 6.25x10^4^ cells/cm^2^ on 13 mm coverslips. 24 hours later cells were incubated with 10 µM CellROX® Deep Red (Life Technologies) and 2.8 µg/ml Hoechst 33342 (Fluka) for 30 minutes. CellROX® and Hoechst 33342 were removed and cells were washed three times with complete medium. Proxison (10 µM) or DMSO vehicle was added for 30 minutes, and then 400 µM *t*BHP was added. One hour after *t*BHP addition, the coverslips were placed into an Attofluor® Cell Chamber (Life Technologies) holder and washed with Hank’s balanced salt solution (HBSS) with calcium and magnesium. Images were taken on the Zeiss Observer of phase contrast, CellROX® Deep Red and Hoechst 33342 fluorescence. Random areas (n=5-6) of each coverslip were imaged. Post-acquisition analysis was performed using Volocity. A small area around each apparent cell nuclei was selected in all images and the mean fluorescence in arbitrary units (au) in these areas was calculated. The mean and standard error of mean (SEM) was calculated (n=4).

### Immunocytochemistry and quantification

SH-SY5Y cells were seeded at 1.25x10^5^ cells/cm^2^ on coverslips and incubated for 24 hours prior to addition of Proxison or vehicle (DMSO) for 30 minutes. Then 400 µM *t*BHP was added for 4.5 hours when cells were fixed with 4% PFA for 20 minutes and washed three times with PBS. Immunocytochemistry was performed by the addition of blocking buffer (0.1% Triton X-100 or 0.3% Triton-X100 for nuclear staining, 2% goat serum, PBS) for 30 minutes. Primary antibodies rabbit IgG NRF2 (1:800, C-20, Santa Cruz), mouse IgG2b 8-OHdG (1:1000, 15A3, abcam) in blocking buffer were added to cells for 16 hours at 4°C, when they were washed 3 times in PBS with 0.1% Triton X-100. Secondary antibodies (Life Technologies) in blocking buffer were incubated with cells for 2 hours at room temperature in foil, when they were washed 3 times in PBS with 0.1% Triton X-100. DAPI (10-50 µg/ml, Life Technologies) in PBS was added for 10 minutes. Cells were imaged using Leica TCS SPE confocal microscope. NRF2 images were quantified using ImageJ. The z-stack images were flattened and a mask was created using the DAPI channel. This mask was then used to calculate the nuclear red fluorescence intensity. The mean fluorescence from each image (2-4 images) and from each coverslip was calculated. The 8-OHdG images were quantified using ImageJ. The z-stack images were flattened and a mask was created using the DAPI channel, this mask included the nuclei and the area surrounding the nuclei by reducing the threshold. This mask was then used to calculate the nuclear green fluorescence intensity. The mean fluorescence from each image was calculated with ImageJ.

### RT-qPCR

SH-SY5Y cells were loaded with Proxison or DMSO as a control for 30 minutes before 400 µM *t*BHP was added for 4.5 hours when cells were lifted with trypsin and pelleted. RNA extraction was performed using the Epicentre MasterPure™ Complete DNA and RNA Purification Kit (Epicentre, MC85200), according to manufacturer’s instructions. RNA concentration was quantified using a Nanodrop spectrophotometer. Total RNA (1 µg) was used for cDNA synthesis. RNase-free water was added to the RNA to give a 10 µl sample. The samples were incubated with 1 µl dNTP mix (10 mM Life Technologies) and 1 µl random primers (50 ng/µl, Thermo Fisher) at 65°C for 5 minutes and then chilled on ice. After brief centrifugation, 4 µl 5x First strand buffer (Life Technologies), 2 µl 0.1M DTT (Life Technologies) and 1µl RNAse OUT (40 units/µl, Life Technologies) were added. The contents was incubated at 37°C for 2 minutes. Then 1 µl M-MLV reverse transcriptase (200 units/µl, Life Technologies) or 1 µl RNase-free water for the negative reverse transcriptase sample was added. This was mixed and incubated at room temperature for 10 minutes when it was moved to 37°C for 60 minutes. The reaction was inactivated by incubation at 90°C for 10 minutes. The cDNA mix was placed on ice and 80 µl RNase-free water was added. The cDNA samples were then ready for RT-qPCR. qPCR was performed using the Roche LightCycler® 480 System with the Universal Probe Library (UPL) (Roche). The Roche UPL Assay design centre was used to design intron-spanning primers with a specific UPL probe for each gene (*TBP* F-gaacatcatggatcagaacaaca R-atagggattccgggagtcat Probe 87; *NRF2* F-acacggtccacagctcatc R-tgcctccaaagtatgtcaatca Probe 18; *HMOX1* F-cagtcaggcagagggtgatag R-agctcctgcaactcctcaaa Probe 42; *NQO1* F-acgctgccatgtatgacaaa R-ggatcccttgcagagagtaca Probe 9). Reactions (10 µl) containing primers, UPL Probe, LightCycle® 480 Probes Master mix (Roche) and PCR water were performed in 386-well plates as described in the manufacturer’s instructions. The results were normalized to RNA levels of TATA-binding protein (*TBP*).

### Treatment of zebrafish embryos with 6-hydroxydopamine (6-OHDA) and neuroprotective compounds

All zebrafish were maintained under Home Office Project Licence No.: 60/4196 and all experiments were performed in accordance with the relevant guidelines and regulations, and approved by the Animal Welfare and Ethical Review Body of the University of Edinburgh. The Wik wild-type strain of zebrafish was used for this study, and they were maintained in standard conditions^54^. Clutches of embryos were obtained from single pair matings to ensure all embryos were at the same developmental stage across treatments. At 24 hours post fertilisation (hpf) embryos were split into groups of 24-30 embryos and treated with 250 µM 6-OHDA (162957, Sigma) or 250 µM 6-OHDA + 10 µM Proxison or quercetin in 3 ml embryo medium for 24 hours (water for control). At 48 hpf embryos were fixed in 4% paraformaldehyde in PBS for 45min. Samples were then blocked (2% bovine serum in 0.1% PBS-triton) for 1h at room temperature. A mouse monoclonal anti-TH antibody (1:1000 diluted in blocking buffer, MAB318, Millipore) was used as the primary antibody and incubated with the samples overnight at 4°C. The next day samples were washed in 0.1% PBS-T, followed by incubation with a secondary antibody Cy3 donkey α-mouse (1:200 diluted in blocking buffer, 715165150, Jackson Immuno) overnight at 4°C. Samples were then washed and mounted in 70% glycerol in PBS. Embryos were imaged by confocal microscopy (LSM710, Zeiss, Oberkochen, Germany), and Z-stacks were taken through the head ensuring all cells could be quantified. Cell counts were performed using ImageJ. The observer was blinded for all quantifications of confocal images.

### Statistical Analysis

Statistical Analysis was performed in Minitab using the Analysis of Variance (ANOVA) statistical test with a Tukey’s post-hoc test.

## Acknowledgements

This work was funded by a Scottish Universities Life Sciences Alliance (SULSA) BioSkape Industry PhD studentship to NJD and Antoxis Limited as the industrial sponsor. TK was funded by a Parkinson’s UK Senior Research Fellowship (F-0902) and an MRC grant (MR/J012831/1). JEL was funded by a Royal Society University Fellowship and grant (RF 120645). MRM was funded by a British Heart Foundation Special Project Grant (SP/15/8/31575). NOD was funded by an MRC PhD studentship. TB and CGB were funded by a BBSRC project grant (BB/M003892/1). We are grateful to Dr Pleasantine Mill and Dr Ammar Natalwala for critical comments on the manuscript. We thank Dr Judith Sleeman for use of cell culture facilities at St Andrew’s University.

### Statement of Author Contributions

TK designed and coordinated the study and wrote the paper. NJD designed and performed cell culture and electroparamagnetic resonance (EPR) experiments. NOD, TB, and CGB designed and performed zebrafish experiments. JEL and MRM designed, supervised, and performed EPR experiments and data analysis. GC and DBM designed experiments and provided Proxison and expertise related to the compound. All authors contributed to and approved the final manuscript.

### Conflicts of Interest

Donald McPhail and Graeme Cook retain shares in Antoxis Limited. The other authors declare no competing financial interests.

